# Ethology as a physical science

**DOI:** 10.1101/220855

**Authors:** André EX Brown, Benjamin de Bivort

**Affiliations:** MRC London Institute of Medical Sciences, London, UK; Institute of Clinical Sciences, Imperial College London, London, UK; Department of Organismic and Evolutionary Biology, Harvard University, Cambridge, MA USA; Center for Brain Science, Harvard University, Cambridge, MA USA

**Keywords:** Behaviour, ethology, data, posture dynamics, discretisation, hierarchy, stereotypy, low dimensionality

## Abstract

Behaviour is the ultimate output of an animal’s nervous system and choosing the right action at the right time can be critical for survival. The study of the organisation of behaviour in its natural context, ethology, has historically been a primarily qualitative science. A quantitative theory of behaviour would advance research in neuroscience as well as ecology and evolution. However, animal posture typically has many degrees of freedom and behavioural dynamics vary on timescales ranging from milliseconds to years, presenting both technical and conceptual challenges. Here we review 1) advances in imaging and computer vision that are making it possible to capture increasingly complete records of animal motion and 2) new approaches to understanding the resulting behavioural data sets. With the right analytical approaches, these data are allowing researchers to revisit longstanding questions about the structure and organisation of animal behaviour and to put unifying principles on a quantitative footing. Contributions from both experimentalists and theorists are leading to the emergence of a physics of behaviour and the prospect of discovering laws and developing theories with broad applicability. We believe that there now exists an opportunity to develop theories of behaviour which can be tested using these data sets leading to a deeper understanding of how and why animals behave.

## Introduction

Animals are material things living in a material world governed by physical laws. Research fields where the influence of physics on behaviour is clearest (such as biomechanics, the function of sensory systems, and the physiology of nervous tissues) have well-developed collaboration between biologists and physicists. This is research aimed at the *physical underpinnings* of behaviour. However, we believe that behaviour itself, at the level of the organism, can be fruitfully studied from a physics perspective. The principled, quantitative approach to animal dynamics represents a nascent *physics of ethology* that is being explored by both experimentalists and theorists.

The conceptual basis for the study of animal behaviour was laid by the first ethologists in the early twentieth century, who conceived their work as *Tierpsychology* (animal psychology)^1^. They were fascinated by the bewildering variety of behaviours in nature and wanted to systematically characterise this variety and to understand why animals behaved in the way they did. A modern variant of this approach is the study of an animal’s location over time, a field sometimes called movement ecology^2^, which has led to fascinating advances on efficient search strategies^3,4^ and the study of collective behaviour^5–7^. However, what interested the early ethologists, and our focus in this article, is the study of behavioural repertoires that include not just where an animal is in its environment, but, in Tinbergen’s words, the “total movements made by the intact animal”^8^. In early ethology, behaviour was typically categorised by trained observers taking advantage of their experience and intuition to identify relevant elements such as feeding, fighting, fleeing, or mating, but Tinbergen’s definition leaves open the question of how behaviour should be measured and represented.

Similar problems arise in physics, where we often have the freedom to measure a vast number of quantities and where the best representation is not always clear. For example, the development of a microscopic theory of superconductivity (BCS) relied not just on experimental observations but also a body of phenomenological theories that each captured an aspect of the problem from a different perspective^9^. By analogy, we believe that alternative representations of animal movement will reveal different phenomena and perhaps inspire new theory. This approach is well-suited to physicists, and its findings will be helpful in identifying lower level mechanisms underpinning behaviour.

To make use of a given representation, the relevant variables must be measurable in an experiment, preferably in large quantities at low cost. Improved cameras and computer vision algorithms have greatly expanded the range of accessible spatial and temporal scales for behavioural quantification as well as the range of environments in which recording is possible. We begin by reviewing how current experimental techniques enable the quantification of behaviour and then consider how measurements are revealing principles of behaviour.

## Techniques

### Posture tracking

Behavioural repertoires are composed of posture changes over time. Quantifying the posture of an organism typically begins with an imaging experiment that records a direct representation of the animal, such as an image of it in a video. From this input, increasingly abstract representations of the animal can be computed using a variety of approaches (Figure 1). Historically, this required the mind-numbing manual annotation of individual video frames. This meant postural quantification for long videos at high framerates was essentially impossible. Stopgap techniques drew inspiration from human motion capture techniques, and placed small dye spots on limbs which, by virtue of their contrast or fluorescence, could be isolated from the image background and automatically tracked^10–12^. Gluing on tracking markers introduces potential confounds, particularly the continuous, unnatural mechanosensory stimulation of subject animals.

**Figure 1.**
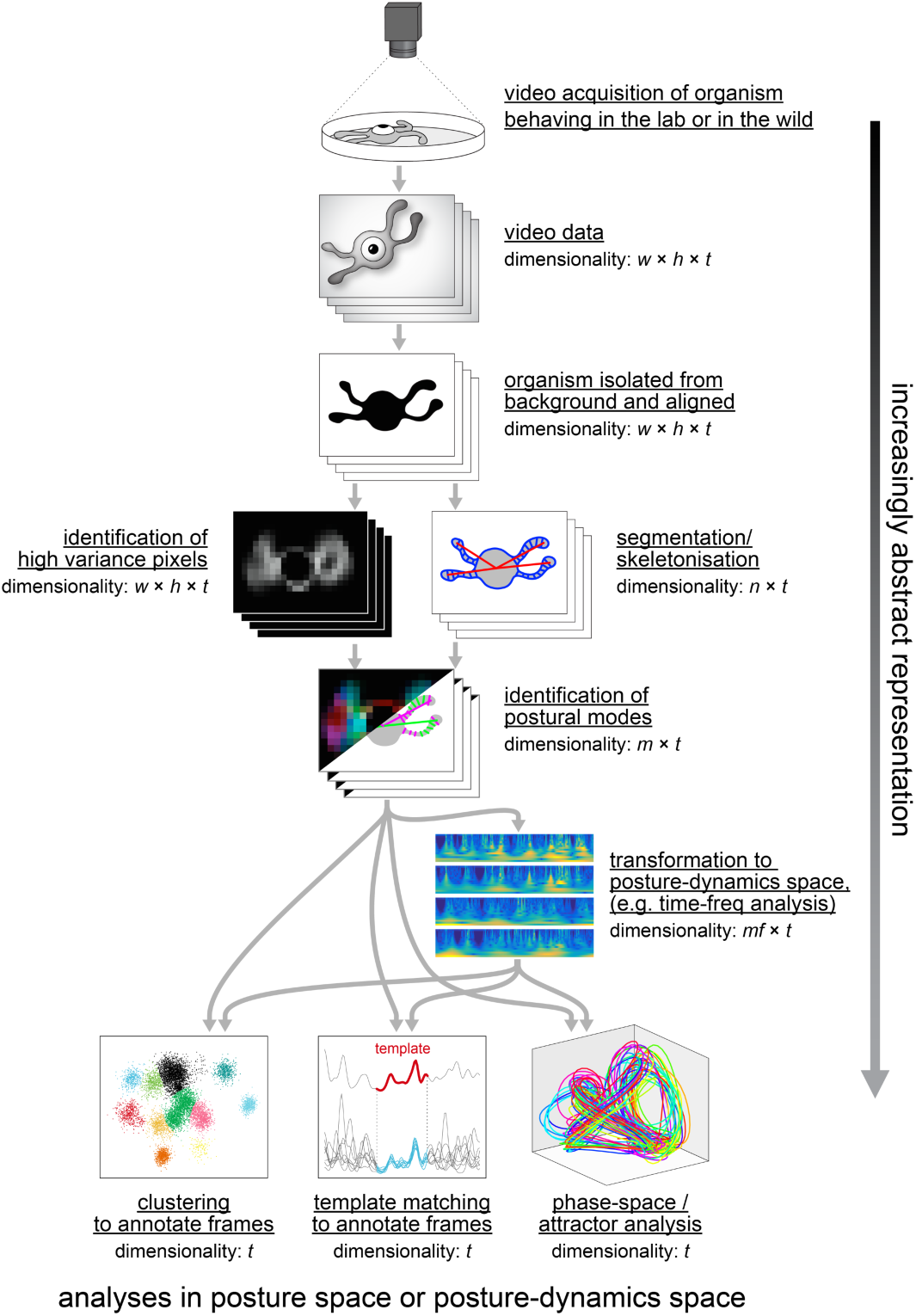
A modern pipeline for the analysis of behavioural data. From a raw video recording, machine vision techniques are used to produce a representation of the animal’s posture in each frame. This can take the form of coordinates of skeletonised landmarks or modes of pixel intensities. This representation can be analysed directly by the techniques shown at the bottom of the pipeline, or first transformed into a posture-dynamics space using, e.g., time-frequency analysis, and then analysed. *w* = width of video frame, *h* = height of video frame, *t* = length of video recording in frames, *n* = number of skeletal landmarks quantified, *m* = number of image or skeletal element modes identified, *f* = number of frequency bands. Generally in such analyses *m* ≈ *f* << *w* ≈ *h* << *t*.

Markerless techniques emerged exploiting frustrated internal reflection to characterise the contact points between an animal and its substrate^13–15^, however these approaches cannot track portions of the animal separated from the substrate. A potential way forward to automatically identify postural landmarks lies in model-based image analysis, in which the known biomechanical degrees of freedom of an animal are used to find the posture that would produce a given image with the highest likelihood^16–18^. The challenges of automatic annotation of subtle behavioural events may be solved by deep learning/ neural network approaches^19,20^. However, as these require human-annotated training sets, the power of the technique may be offset by the inconsistency in human labels, which vary from lab to lab and expert to expert^21,22^.

An alternative to using image analysis to extract landmark-based or skeletonised representations is to use the raw pixel luminosities in animal-aligned video frames (Figure 1). Identification of the principal components (eigenvectors of the covariance matrix) of pixel luminosity reveals the regions of the animal that change during behaviour^23,24^. The advantage of this pixel-based representation is that it can be applied to animals of any shape or limb configuration.

### Real-time feedback and brain imaging

The methods described above vary greatly in their computational intensity. Some computationally lighter methods can be performed in real time, like fitting an ellipse to the silhouette of an adult *Drosophila melanogaster* fly^25^ or segmenting the midline of a *Caenorhabditis elegans* round worm^26^. This enables closed-loop experiments in which the animal is stimulated (either through its sensory systems^25,27^ or directly in the nervous system using optogenetics^25,27–29^) in response to behaviours it performs in real time, as soon as they are flagged. This also permits real-time prediction of upcoming behaviours, which can facilitate close tracking of animals, improving the quality of neural activity recordings acquired from freely moving animals^30^, which faces substantial challenges both in tracking individual animals for long periods and in registering volumetric neural imaging data.

### Bridging behavioural timescales

A common feature of long behavioural recordings is modulation on multiple timescales. The swim bouts of zebrafish last a few tens of milliseconds^31^, and flies can turn in response to odour stimuli within 70ms, less than the length of a single running stride^32^. And yet, the behavioural repertoire of both of these organisms is tuned by long timescale processes such as satiety/ hunger^33^ and the circadian cycle^25^. Behaviour changes over the longest timescale possible in organisms, the lifetime^34^, and quantitative variation in behaviour with age is conspicuous in many measurements.

While it remains challenging to acquire high resolution video data over long time periods, there are approaches that permit the measurement of behaviour on timescales of lifetimes. Such longitudinal experiments are naturally most advanced for organisms with short lifespans, such as single-celled microbes where tracking of entire cohorts for multiple generations is possible^35^. Long-term recordings are facilitated by real-time processing of video data to extract and save only compressed representations of behaviour, such as animals’ centroids. Such low dimensional representations facilitate not just experiments in depth (over time), but also in breadth (over individuals). As an example, tracking the centroids of many flies in parallel exploring simple arenas has permitted the measurement of individual behaviour from tens of thousands of individuals across hours and days^36,37^. In worms, midline tracking can be done in real time for dozens of individuals^26^. Combining multi-worm midline tracking with custom sample chambers^34,38^ would enable cradle-to-grave tracking of an animal’s complete movements with sub-second resolution, providing valuable information about behavioural ageing and healthspan that could be combined with genetic and drug perturbation and compared with other physiological changes. In mice, the highest resolution tracking is typically done for limited periods in fairly featureless ‘open fields’^39,40^, but longer timescales can be accessible in systems that track mice in their home cages^41–44^.

### Inexpensive and field imaging

At the same time that new instrumentation has permitted deep and broad behavioural recordings, the affordability and versatility of behaviour rigs has improved. There are now open source designs for behaviour recording devices that capture behaviour from many individuals simultaneously while also permitting closed-loop stimulation of animals^25,45^. These technologies are flourishing in part because of the proliferation of inexpensive hardware for data acquisition, such as single-board computers (e.g., Raspberry Pi) and convenient break-out microcontroller boards (e.g., Ardunio and Teensy). This open hardware community thrives in the groundwork laid by earlier open source animal tracking algorithms^46–48^. We believe there is great potential to bring these technologies out of the lab and into the field to capture behaviour in circumstances closer to that in which it evolved, its proper ethological context^49^.

## Principles

The growing technical sophistication of behavioural studies has correlated with growing conceptual insights. Several principles have been identified to help guide exploration of animal behaviour (Figure 2). Some of the principles have close parallels in traditional physics, while others appear to be uniquely biological. Some are more tentative, while others have been found in diverse species in diverse contexts. Overall, they highlight regularities in behaviour that may lead to novel theory.

**Figure 2.**
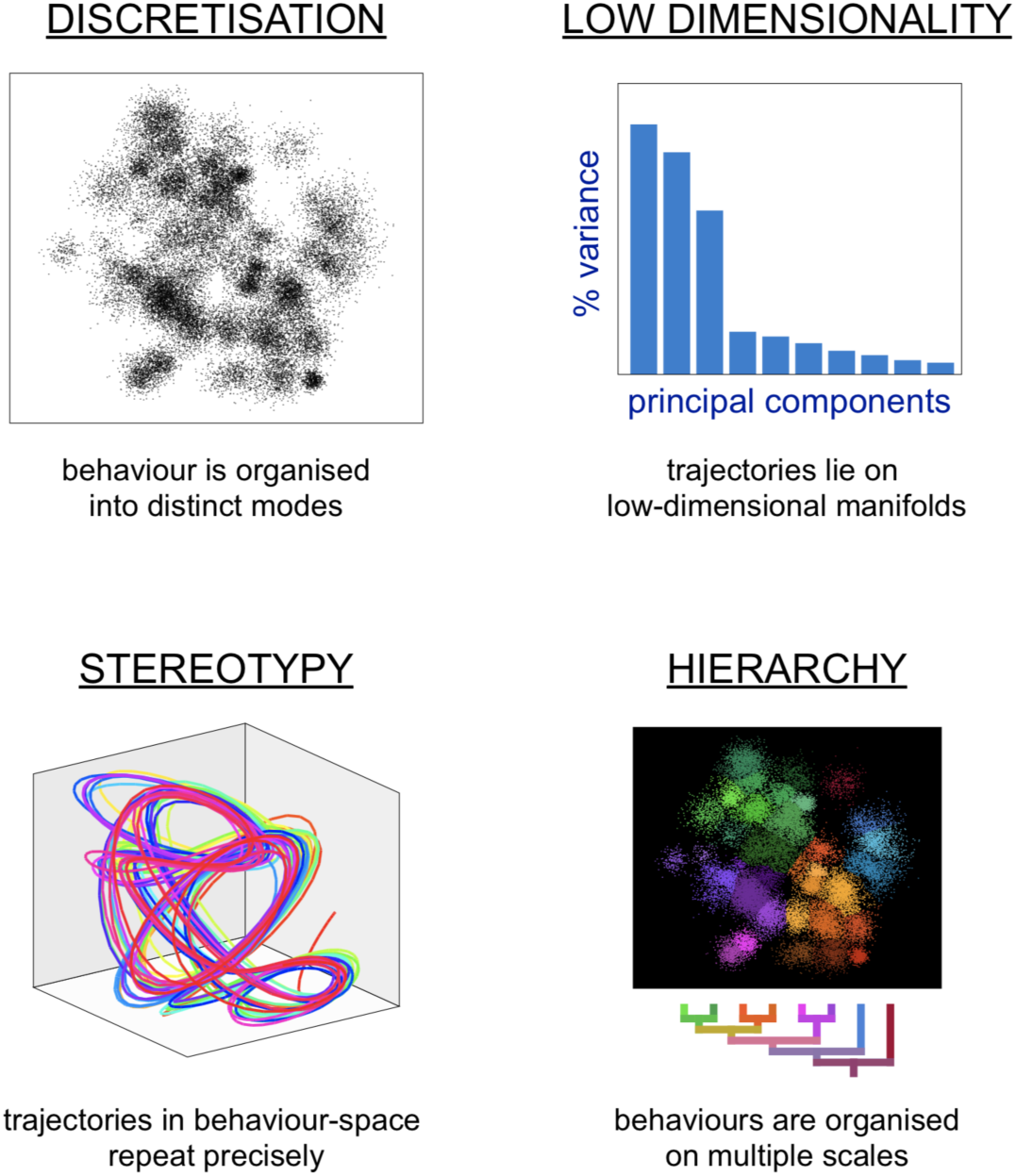
Principles of the organisation of behaviour. Analyses of behaviour in multiple species using multiple approaches have pointed to four principles organising behaviour. Discretisation: behaviour occurs in distinct patterns that appear as clusters in representations of behaviour space. Low dimensionality: behaviour can be specified with fewer values than the morphological degrees of freedom available to the animal. Stereotypy: instances of behaviour within a distinct cluster can be strikingly similar in implementation. Hierarchy: behaviour is organised on multiple scales, and is naturally categorisable into behavioral states with sub-states (and sub-sub-states, etc.).

### Low postural dimensionality

In principle, animals have many postural degrees of freedom. However, during normal behaviour, correlations between body parts make it possible to derive lower dimensional representations of posture that reveal aspects of coordination and also provide a useful basis for other behavioural analyses. In *C. elegans*, the length-wise midline predicts with high fidelity the position of the entire body. This was exploited to determine the dimensionality of worm posture during spontaneous exploration and it was found that four dimensions explain 95% of the postural variance^50^. Two of these “eigenworms” correspond approximately to a quadrature pair that supports travelling waves with a third mode capturing overall curvature. Related methods agree on the dimensionality but highlight different aspects of locomotion^51,52^. Scrutiny of dynamics in worm postural space showed that direction changes can be modelled as stochastic switches between attractors^53^ and revealed a new behaviour (the delta turn) that previous investigators had missed^54^. Related analyses have been applied in larval zebrafish where the midline also captures much of the postural information^55^.

Low dimensionality compared with the degree of postural freedom also holds for animals with more complex body plans. Monty Python’s Ministry of Silly Walks is funny and modern dance is striking in part because they explore regions of postural space that lie off the manifold that captures most behavioural dynamics. More constrained tasks such as reaching in humans and monkeys also show low dimensionality, leading to work on muscle synergies and investigations into the neural control of reaching^56,57^. Interestingly, even octopuses, whose flexible limbs allow for higher degrees of freedom seem to create ‘joints’ when reaching, potentially as a way of addressing the otherwise high dimensionality of the task^58^.

That behaviour lies in low dimensional subspaces holds in both posture-spaces and posture-dynamics spaces. In the former, data points represent an animal’s posture at a specific time^50^. In the latter, data points represent posture and its change over time, as computed by time-frequency analysis^23,59^ (Figure 2) or slicing recordings into segments of a particular length^60^. Interestingly, all of the principles of the organisation of behaviour described in this review—low dimensionality, discretisation/stereotypy and hierarchy—emerge in analyses of both posture space and posture-dynamics space.

### Stereotypy and discretisation

Behaviour is often thought of as a sequence of discrete actions that can be represented in an ethogram, a network diagram in which nodes are actions and edges transition rates. An assumption at the heart of this approach is that animals perform some actions in ways that are similar enough to previous instances (and different enough from instances of other actions) to be discretely classifiable. Similarity across instances of an action is called stereotypy and it can be quantified by determining how long it takes for trajectories through posture or posture-dynamics space to diverge. Lyapunov exponents and trajectory correlation time constants serve as metrics of stereotypy. For example, adult fly velocity trajectories remain correlated for a few hundreds of milliseconds^61^.

Stereotypy is the basis for several recent methods for segmenting behaviour into actions without the requirement of human-annotated sequences for training. An advantage of these unsupervised segmentations is that it is possible to discover actions that may have been difficult to detect initially by human observations. In *C. elegans*, behavioural motifs were identified as precisely repeated segments in postural time series and their variation across hundreds of mutants revealed some underlying genetic relationships^60^. In *Drosophila* larvae, onset of optogenetic neuronal stimulation was used to align trajectories and unsupervised structure learning detected distinct behavioural phenotypes^62^. In adult flies, parameterized high resolution video of spontaneous behaviour of adult Drosophila using time-frequency analysis of principal components of video frame pixels^23^. This method has since been extended to pairs of socially interacting animals^24^. In zebrafish larvae, stereotypy was identified by a novel density-based clustering algorithm in a feature space defined by multiple movement and postural parameters^63^.

Unsupervised methods have many methodological choice points and tuning parameters, and navigating these without reverting to supervised analysis is a conceptual challenge. One approach is to favour methods that produce behavioural annotations most consistent with axiomatic assumptions of what behaviour is^59^, and this approach could be extended to choosing methods based on their adherence to physical rules or probabilistic models.

So far, unsupervised approaches have revealed stereotyped patterns that are at least somewhat discretised. That is, non-stereotyped patterns fill the interstices between stereotyped modes in behaviour space^23^. Reusing a finite set of well-adapted actions could be an efficient way of organising a behavioural repertoire, but too much stereotypy comes with costs. For example, when circumstances change, animals are able to adapt flexibly, performing rare or even unique bouts of activity as required. Compare, for example, the high stereotypy of a person walking down an empty corridor with the novel sequences of stops, starts, and skips that is observed when moving against the flow of a crowd. This may reflect a trade-off between efficient stereotyped behaviour and relatively inefficient but necessary flexible behaviour. A similar efficiency argument has been advanced to underlie Zipf’s law^64^, the power-law describing the rank-frequency distributions of word use^65^ and heavy-tailed distributions of behaviours have also been observed in spontaneous worm behaviour^66^. However, in some circumstances at least, heavy-tailedness observed at a population level reflects heterogeneity among animals that are individually stereotyped^67,68^.

Stereotypy is also intimately connected with predictability which can carry its own costs, especially in the presence of predators. Tentacle snakes take advantage of the predictability of escape responses to capture fish^69^. More generally, behavioural variability across or within individuals may reflect bet-hedging strategies to cope with unpredictability in the environment^70^. Neural circuits that increase noise above the constraints imposed by sensory information have been found to actively increase behavioural variability even in seemingly simple sensory pathways^71^. In humans, higher task variability is associated with faster motor learning^72^.

### Hierarchical organisation

Much like a language organised into phrases, sentences, paragraphs, and documents, hierarchical organisation has been proposed as a guiding principle in the analysis of behaviour^8,73^. It has been shown that hierarchical organisation can be an efficient means of using limited computation to produce complex adaptive behaviour in robotics^74^, and the example of language has long been an important inspiration in the study of sequential behaviour^75^, but clear demonstrations of hierarchical organisation in non-human animals have been lacking.

The simplest form of hierarchy is a linear ‘peck order’ in which higher level behaviours dominate lower level behaviours until they are complete. Such a suppression hierarchy has been recently observed in grooming behaviour in fruit flies^76^. Nested representations with multiple levels of nesting have been constructed from action sequences using dictionary-based compression algorithms in worms^77^ and cluster analysis in fish^63^. However, results from intrinsically hierarchical algorithms do not necessarily imply that behaviour is organised hierarchically. A notable exception in the hierarchical analysis of behaviour comes from work on adult fruit flies. Starting from the observation that transitions between behavioural states in flies are non-Markovian, Berman *et al.* have used a treeness metric to quantify the degree of hierarchy in flies’ behavioural repertoire^78^ and showed that a hierarchical representation is optimal for predicting the flies future behaviour with progressively coarser-grained representations predicting behaviours at longer time scales.

## Emerging directions

We have focused on methods and principles for studying behavioural organisation. The understanding gained by these approaches is richer still when it reveals relationships between behaviour and genetic variation, neural activity, or evolution. As quantitative behavioural representations are maturing, elucidating relationships with other lower and higher biological levels presents several outstanding questions.

An often-implicit hypothesis is that behavioural features characterised using the methods we have reviewed will fruitfully map onto other variables of interest, most prominently genetic variation and neural activity. However, this is an empirical hypothesis not guaranteed by theory. An exciting current area of research is in defining behavioural representations jointly with other variables of interest. For example, behaviours that covary with genetic variants in populations of organisms^79,80^ could provide better mappings between genotype and phenotype. Rather than checking for such relationships with a behavioural representation already set in stone, it may be possible to develop algorithms (or new objective functions for existing algorithms) that simultaneously optimize a behavioural representation and its mapping to complementary genetic data.

Similarly, the relationships between neural circuits or patterns of activity in large populations of neurons and behaviours could be learned by considering their representations simultaneously. The promise of this notion is evident in the rich variations in supervised^81^ and unsupervised^82^ annotations of *Drosophila* behaviour induced by systematic perturbation of circuit elements. The behavioural elements highlighted through such joint analyses may turn out to be quite different from the elements identified in behaviour-only approaches. Conversely, it is likely that joint analyses of behaviour and neural circuit dynamics will provide improved representations and understanding of physiology. That is, neural coding may only make sense through the lens of behaviour^83–85^.

Behaviour is a proximal determinant of animal fitness. Thus it is likely that the organization of behaviour shapes evolutionary trajectories and is in turn shaped by evolution. Fisher’s geometric model argues that high-dimensional phenotypes create a large space for evolution to explore and thus reduce the probability that a random mutation is beneficial^86^. Does the discrete organisation of behaviour imply many degrees of freedom producing such a large space or do genetic correlations between behaviours reduce the effective dimensionality? Does the hierarchical organisation of behaviour constrain this space to make behavioural evolution more efficient?

We have examined how new techniques are producing large data sets that have recently revealed a few high-level principles in the organisation of behaviour. But some of the most interesting directions for future research may be the codification or derivation of these principles from formal theory, the approach which has been so successful in physics. Given the increasingly precise and diverse datasets that are now available, there are ample opportunities to be inspired from behaving animals and to test models quickly. Major open questions focus exclusively on behaviour itself (what are the equations of motion for an organism?) and also bridge levels of biological organisation (How is behaviour jointly distributed with neural activity? How does its organisation constrain evolution?). Ultimately, we hope the impact of physics on ethology will be akin to its impact on molecular biology, where ideas like the genetic code and techniques like X-ray crystallography have been key in understanding life.

## Acknowledgements

BdB was supported by a Sloan Research Fellowship, National Science Foundation grant no. IOS-1557913 and Klingenstein-Simons Fellowship Award in Neuroscience. AEXB was supported by a Medical Research Council grant MC-A658-5TY30. We thank an anonymous reviewer for thoughtful suggestions that expanded the manuscript’s dimensions.

